# MG53 Protects Against Intestinal Inflammation by Inhibiting NLRP3 Inflammasome Activation

**DOI:** 10.64898/2026.02.18.706669

**Authors:** Zhongguang Li, Zach Dawson, Xiangguang Li, Serenali Zhao, Matthew Bu, Fei Jiang, Yuchen Chen, Min Zhang, Xindi Zeng, Ki Ho Park, Jing Lu, Jinshan He, KyungEun Lee, Prosper Boyaka, Haichang Li, Jianjie Ma

## Abstract

**Background and Objectives:** Inflammatory bowel disease (IBD) involves dysregulated immune responses and chronic intestinal inflammation. The nod-like receptor pyrin domain–containing 3 (NLRP3) inflammasome plays a critical role in IBD pathogenesis, but regulatory mechanisms remain not fully understood. Mitsugumin 53 (MG53, also known as TRIM72), originally identified as a critical membrane repair protein, has emerged as a novel regulator of inflammatory processes. To investigate the protective role of MG53 in colitis and elucidate its mechanisms in regulating NLRP3 inflammasome activation in IBD.

**Methods:** We used dextran sodium sulfate (DSS)-induced colitis models comparing *MG53* knockout (*MG53*^−/−^) and wild-type (WT) mice, assessing disease severity, MG53 tissue uptake, and therapeutic effects of recombinant human MG53 (rhMG53). *In vitro* studies examined rhMG53’s effects on NLRP3 inflammasome activation, caspase-1 cleavage, interleukin-1β (IL-1β) secretion, and MG53-NLRP3 interactions.

**Results:** *MG53*^−/−^ mice showed more severe colitis with increased weight loss, higher disease activity scores, shortened colons, and greater inflammation. DSS treatment induced the accumulation of circulating MG53 in inflamed colonic tissue. rhMG53 administration ameliorated colitis severity in *MG53*^−/−^ mice and dose-dependently suppressed NLRP3 inflammasome activation *in vitro*. MG53 interacted with NLRP3 and reduced apoptosis-associated speck-like protein containing a CARD (ASC) speck formation and NLRP3 oligomerization without affecting upstream signaling or NLRP3 stability.

**Conclusion:** MG53 is a physiological regulator of NLRP3 inflammasome activation that protects against colitis, suggesting therapeutic potential for IBD.

## Introduction

Inflammatory bowel disease (IBD) represents a significant global healthcare burden. According to recent epidemiological data, approximately 2 million U.S. adults (1.0 % of the adult population) have been diagnosed with IBD(1, 2). IBD includes a spectrum of chronic, relapsing-remitting inflammatory disorders of the gastrointestinal tract, with Crohn’s disease (CD) and ulcerative colitis (UC) being the two predominant clinical manifestations. The pathogenesis and molecular underpinnings of IBD remain incompletely understood - accumulating evidence identifies multifactorial dysregulation of mucosal immune responses, compromised intestinal barrier function, and aberrant innate immune signaling as key biological drivers. IBD significantly impairs health-related quality of life, increases long-term disability, and confers elevated risk of colorectal malignancy one of the most prevalent cancers in adults(3, 4). Despite significant advances in understanding IBD pathophysiology, current therapeutic options remain limited, highlighting the urgent need for novel therapeutic targets and approaches(5-7).

The innate immune system plays a pivotal role in intestinal pathogenesis, with pattern recognition receptors serving as key mediators of inflammatory responses. Among these, the nod-like receptor pyrin domain–containing 3 (NLRP3) inflammasome has emerged as a critical component in intestinal inflammation(8, 9). The NLRP3 inflammasome is a multimeric protein complex consisting of the NLRP3 sensor, the adaptor protein apoptosis-associated speck-like protein containing a CARD (ASC), and the effector caspase-1(8). Upon activation by diverse stimuli including pathogen-associated molecular patterns and damage-associated molecular patterns, NLRP3 undergoes oligomerization and recruits ASC to form large supramolecular structures called ASC specks(10). This assembly leads to caspase-1 activation and subsequent cleavage of pro-IL-1β and pro-IL-18 into their mature, biologically active forms, which are then secreted to propagate inflammatory responses.

Accumulating evidence demonstrates that NLRP3 inflammasome activation is strongly associated with IBD development and progression. Elevated expression of NLRP3 components and increased IL-1β production have been consistently observed in intestinal tissues from IBD patients and animal models. The inflammasome is widely expressed in both intestinal epithelial cells and immune cells, including macrophages, dendritic cells, and neutrophils, making it a central hub for coordinating inflammatory responses in the gut. However, while NLRP3 inflammasome activation contributes to host defense against pathogens, excessive or prolonged activation can lead to chronic inflammation and tissue damage characteristic of IBD(11). Given the critical role of NLRP3 inflammasome in IBD pathogenesis, understanding the regulatory mechanisms that control its activation is essential for developing targeted therapeutic strategies(11-13). Various endogenous negative regulators have been identified, but the complete regulatory network remains incompletely characterized(14, 15).

MG53, a novel tripartite motif-containing (TRIM) family protein, has been identified as a critical effector of cellular membrane repair mechanisms(16, 17). Extensive investigations have elucidated its biological functions in tissue repair processes and therapeutic applications in regenerative medicine (17-29). *MG53* knockout (*MG53*^−/−^) mice exhibit multisystem pathology affecting skeletal muscle, cardiac tissue, and integumentary structures, attributed to impaired cellular membrane repair capacity. Conversely, transgenic mice engineered for systemic MG53 overexpression maintain normal lifespan while demonstrating enhanced tissue repair kinetics and regenerative responses following injury(29, 30). We and other investigators have established that recombinant human MG53 (rhMG53) confers cyto-protection against membrane damage across diverse cell lineages and ameliorates pathophysiology in preclinical models of muscular dystrophy(16, 19, 31), acute lung injury(19, 22, 32), myocardial infarction (33), acute and chronic kidney injury(21), and wound healing(20, 34). Moreover, parallel studies have characterized MG53 as an E3 ubiquitin ligase that targets insulin receptor (IR) and insulin receptor substrate-1 (IRS-1) for proteasomal degradation, thereby contributing to insulin resistance pathogenesis(26, 35, 36). Beyond its established role in tissue repair, our recent investigations have revealed that MG53 exhibits anti-inflammatory properties and provides protection against influenza virus-induced acute lung injury(27, 37).

The potential connection between MG53 and inflammasome regulation has not been explored in the context of IBD. Given the critical role of NLRP3 inflammasome in intestinal inflammation and the emerging anti-inflammatory properties of MG53, we hypothesized that MG53 regulates NLRP3 inflammasome activation and contributes to IBD pathogenesis. In this study, we investigated the role of MG53 in experimental colitis and its regulatory effects on NLRP3 inflammasome activation. Using *MG53* knockout mice and the dextran sulfate sodium (DSS)-induced colitis model(38, 39), we demonstrate that *MG53* deficiency exacerbates intestinal inflammation, while rhMG53 treatment protects against DSS-induced disease pathologies. Using bone marrow-derived macrophages (BMDMs) and human macrophage cell lines, we show MG53 interacts with NLRP3 and serves as a negative regulator of inflammasome activation by inhibiting NLRP3 oligomerization and ASC speck formation. Together, our findings identify MG53 as a novel therapeutic adjuvent for IBD treatment and provide new insights into the regulatory mechanisms controlling intestinal inflammation.

## Materials and Methods

### Regents, Antibodies, rhMG53

Lipopolysaccharide (LPS) from *Escherichia coli* 0111: B4 (tlrl-eblps), Nigericin sodium salt (tlrl-nig), adenosine 5′-triphosphate disodium salt (ATP, tlrl-atpl) and monosodium urate crystals (MSU, tlrl-msu) were from InvivoGen. MG132 (474787), Phorbol 12-myristate 13-acetate (PMA, 79346) and disuccinimidyl suberate (S1885-1G) were obtained from Sigma Aldrich. DSS (160110) was from MP Biomedicals. Anti-MG53 is a custom-generated rabbit polyclonal antibody (16). Anti-NLRP3 (AG-20B-0014), anti-Caspase-1 (AG-20B-0042) and anti-ASC (AG-25B-0006-C100) were from AdipoGen. Anti-CD11b (ab133357) and anti-IL-1β (ab9722) were from Abcam. Anti-IL-18 (54943), anti-β-Actin (4970), anti-GAPDH (2118), HRP-labeled Goat Anti-Rabbit IgG (7074S), HRP-labeled Goat Anti-Mouse IgG (7076S) and HRP-labeled Mouse Anti-rabbit IgG (Light-Chain Specific, 45262) were from Cell Signaling Technology. The rhMG53 was purified as described previously(19). The rhMG53 was stored as lyophilized powder and dissolved in saline solution before use.

### Animals Care and in vivo IBD Mouse Model

All animal experiments were conducted in accordance with the guidelines of the National Institutes of Health (NIH) and were approved by The Ohio State University Institutional Animal Care and Use Committee (IACUC). Mice were maintained in the barrier facility of The Ohio State University Laboratory Animal Resources under controlled temperature, humidity, and a 12 h light/dark cycle. C57BL/6 WT mice were obtained from Jackson Laboratory. *MG53*^−/−^ mice and *MG53*-overexpressing transgenic mice (TPA) were bred and generated as previously reported(29, 40).

For DSS-induced colitis(38, 41), ten-to twelve-week-old male mice were used. Colitis was induced by providing 3% (w/v) DSS in drinking water for 9 consecutive days. For the experiment on the therapeutic efficacy of rhMG53, DSS (3%, w/v) was administered in drinking water for 5 consecutive days, followed by replacement with regular drinking water. Recombinant human MG53 (rhMG53, 2 mg/ml) or vehicle intraperitoneally injected daily from day 5 to day 10. Body weight, stool consistency, and fecal blood were monitored daily. Disease activity index (DAI) was scored based on weight loss, stool consistency, and bleeding according to established criteria. At the end of the experiment, mice were euthanized, and colons were collected, measured for length, and either snap-frozen in liquid nitrogen for protein analysis or fixed in 4% paraformaldehyde for histological and immunofluorescence analysis. Histological scoring was performed on hematoxylin and eosin (H&E)-stained colon sections collected on day 9 of DSS treatment, following previously published scoring systems(42).

### Cell Culture and Plasmid Transfection

THP-1, L929, and HEK293T cells were obtained from ATCC and maintained at 37 °C in a humidified 5% CO_2_ incubator. THP-1 cells were cultured in RPMI-1640 medium, whereas L929 and HEK293T cells were cultured in DMEM. All media were supplemented with 10% fetal bovine serum, penicillin (100 U/ml) and streptomycin (100 μg/ml). For plasmid transfection, HEK293T was transfected using Lipofectamine™ 3000 (Thermo Fisher, L3000015) following the manufacturer’s protocol. The pEGFP-C2-NLRP3 plasmid (73955) was purchased from Addgene. The pmR-mCherry-MG53 plasmid was constructed in our laboratory as previously described(43). After 24-48 h, cells were harvested for subsequent analysis.

### Isolation and Culture of Mouse BMDMs

BMDMs were obtained from 8–12-week-old mice. After euthanasia and sterilization with 75% ethanol, femurs and tibias were dissected and bone marrow was flushed with serum-free DMEM. Cells were collected, subjected to red blood cell lysis (BioLegend, 420301), filtered through a 70 μm strainer, and centrifuged. The resulting cells were cultured in complete DMEM supplemented with 20% L929 - conditioned media derived from a density of 1 × 10^6^ cells/ml. After 7 days of culture with medium change on day 3, mature BMDMs were used for subsequent experiments.

### Activation of NLRP3 Inflammasome in Macrophages

Differentiated THP-1 cells by PMA (100 nM, 24 hours) were plated in 12-well plates and primed with 0.5 μg/ml LPS for 3 hours. BMDMs were plated in 12-well plates and cultured overnight, then primed with 100 ng/ml LPS in Opti-MEM (Thermo, 31985070) for 3 h. Following LPS priming, cells were treated with rhMG53 at 1, 5, or 10μg/ml for 1 h, and subsequently stimulated with either 5mM ATP for 30 min, 10μg/ml nigericin for 1 h, or 150μg/ml MSU for 6 h. After stimulation, cells and culture supernatants were collected. Supernatants were centrifuged at 1,000 rpm to remove debris, and clarified supernatants were snap-frozen in liquid nitrogen and stored at −80°C for subsequent assays.

### ASC Speck Preparation and Detection

After treatment, cells were washed three times with Phosphate-Buffered Saline (PBS) and lysed on ice for 10 min in Triton buffer [50 mM Tris-HCl (pH 7.5), 150 mM NaCl, 0.5% Triton X-100, EDTA-free protease inhibitor cocktail (Sigma, P8340-5ML). Lysates were centrifuged at 6000 × g for 15 min at 4°C, and the supernatant was collected and mixed 1:1 with 2× SDS-PAGE loading buffer, followed by heating at 95 °C for 10 min. The pellet was washed once with Triton buffer, centrifuged again under the same conditions, and then resuspended in Triton buffer containing 2 mM Disuccinimidyl suberate. Crosslinking was carried out at room temperature with gentle rotation for 30 min. After centrifugation (6000 × g, 15 min, 4°C), the pellet was resuspended in 50 μl of 1× SDS-PAGE loading buffer and subjected to 12% SDS-PAGE for analysis.

### Measurement of IL-1β Secretion by Enzyme-Linked Immunosorbent Assay (ELISA)

The secretion of IL-1β from stimulated THP-1 cells or BMDMs was measured using human or mouse IL-1β ELISA kits (R&D Systems, DY201 for human, DY401 for mouse) following the manufacturer’s instructions. Briefly, cell culture supernatants were collected after inflammasome activation, centrifuged at 1,000 × g for 5 min to remove cellular debris, and the clarified supernatants were either used immediately or stored at −80 °C until assay. Standards and samples were added to the ELISA plate wells, followed by the sequential addition of detection antibodies and substrate solution. The absorbance was measured at 450 nm using a microplate reader, and IL-1β concentrations were calculated based on the standard curve.

### Western Blotting and Co-immunoprecipitation (Co-IP)

Cells or colon tissues were lysed in RIPA buffer (150 mM NaCl, 10 mM Tris-HCl, pH 7.2, 0.5% SDS, 1% NP-40, 0.5% sodium deoxycholate) supplemented with protease and phosphatase inhibitors. Cells were washed with PBS and lysed on ice for 20 min. Colon tissues were rinsed with cold PBS, cut into small pieces, and homogenized on ice in RIPA buffer. Lysates were centrifuged at 15,000 × g for 10 min at 4°C, and the supernatants were collected. Protein concentrations were determined using a BCA assay (Bio-Rad, 5000002). Samples were mixed 1:1 with 2× SDS loading buffer (100 mM Tris-HCl, pH 6.8, 200 mM dithiothreitol, 4% SDS, 0.2% bromophenol blue, 20% glycerol), boiled at 100 °C for 10 min, aliquoted, and stored at −80°C.

Equal amounts of protein were separated by SDS-PAGE and transferred to PVDF membranes (MP, IPVH00010). Membranes were blocked with 5% non-fat milk in TBST (20 mM Tris, 150 mM NaCl, 0.1% Tween-20) at room temperature for 1h, then incubated with primary antibodies diluted in blocking buffer overnight at 4°C. After three washes with TBST (10 min each), membranes were incubated with HRP-conjugated secondary antibodies at room temperature for 1h and washed again three times. Protein bands were visualized using enhanced chemiluminescence (Thermo Scientific, 34095) and quantified with ImageJ software.

Co-IP was performed using the Pierce™ Classic Magnetic IP/Co-IP Kit (Thermo, 88804) following the manufacturer’s instructions. Briefly, HEK293T cells were transfected with the indicated plasmids for 48 h before harvest. 500 μg of cell lysate was incubated overnight at 4°C with 5 μg of the indicated antibody (anti-MG53, anti-NLRP3, or IgG control). The immune complexes were captured with Protein A/G magnetic beads, washed, and eluted. The eluates were neutralized and analyzed by immunoblotting.

### Histology and Immunofluorescence Staining

Histology and immunofluorescent staining were performed as previously described(20). Treated cells were washed with PBS, fixed with 4% paraformaldehyde for 10 min, permeabilized with 0.5% Triton X-100 for 5 min, and blocked with 3% Bovine Serum Albumin (BSA) for 1 h. Primary antibodies (ASC: 1:100) were incubated overnight at 4°C, followed by Alexa Fluor 488- or 647-conjugated secondary antibodies (1:500) for 1 h at room temperature. Nuclei were stained with DAPI, and coverslips were mounted with antifade medium.

Paraffin-embedded colon sections were deparaffinized, rehydrated, and subjected to antigen retrieval in citrate buffer (pH 6.0). Sections were blocked with 3% BSA for 1 h, incubated with anti-CD11b (1:100) overnight at 4°C, followed by Alexa Fluor-conjugated secondary antibody (1:500) for 1 h. Nuclei were counterstained with DAPI, and sections were mounted with antifade medium. Images were taken by Olympus BX43 microscope and Nikon Ti-2 Eclipse Confocal Fluorescence Microscope.

### Statistical Analysis

The data were analyzed by GraphPad Prism 10.0 software and are presented as mean ± SD. For comparisons between two groups, statistical significance was determined using an unpaired two-tailed Student’s t-test. For multiple group comparisons, two-way ANOVA was used. P values were provided as \**p* < 0.05, \*\**p* < 0.01, and \*\*\**p* < 0.005.

## RESULTS

### Loss of MG53 Exacerbates DSS-induced colitis

To elucidate the function of MG53 in inflammatory bowel disease pathogenesis, age-matched male WT and *MG53*^−/−^ were subjected to 3% DSS treatment, and their colitis phenotypes were analyzed. On day 9 of DSS administration, *MG53*^−/−^ mice displayed markedly more severe colitis compared to WT controls, as evidenced by pronounced body weight loss (*p* < 0.01, **Fig. 1A**), elevated DAI scores (*p* < 0.01, **Fig. 1B**), and significantly shortened colons (*p* < 0.01, **Fig. 1C and D**).

**Figure 1.**
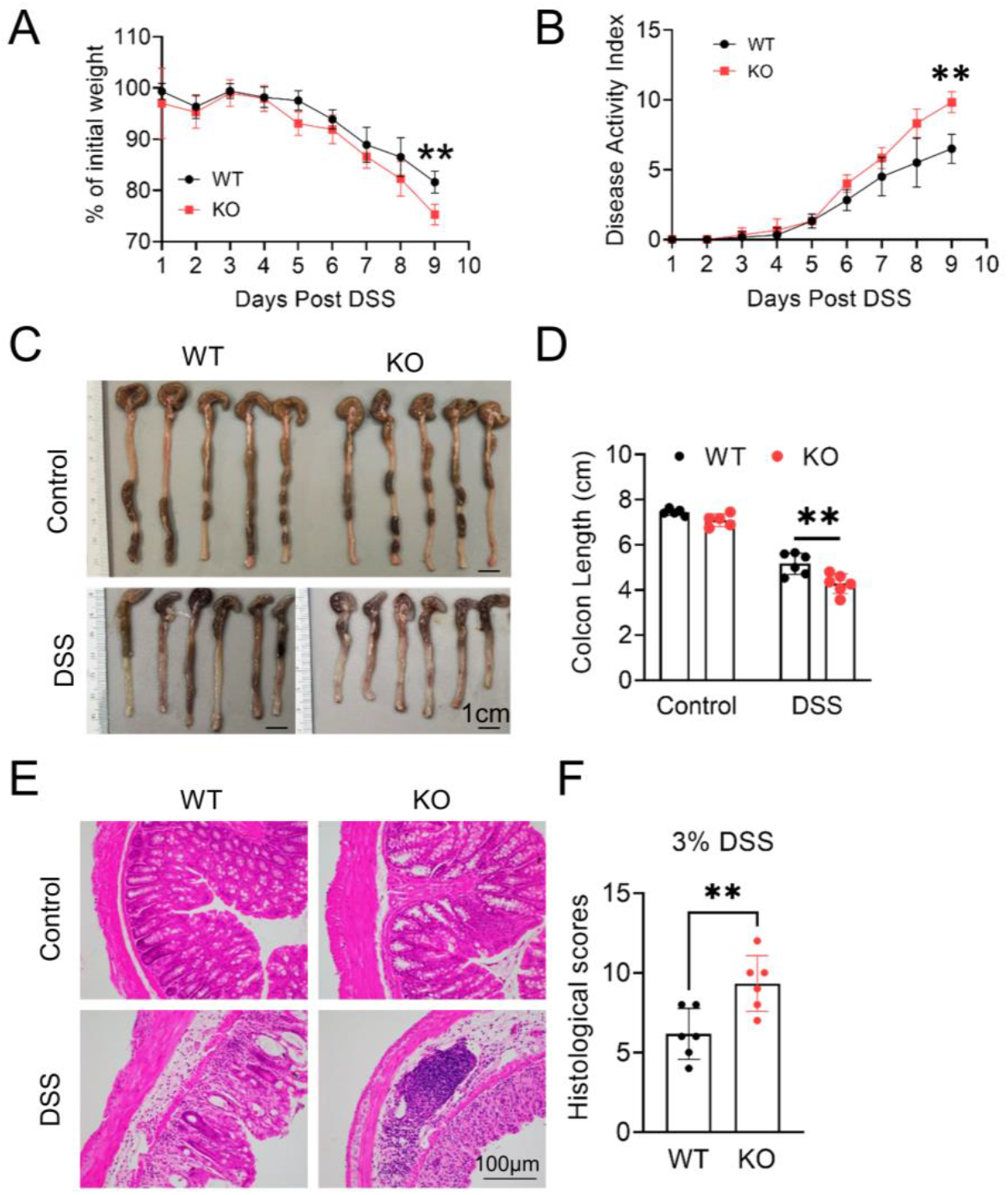
MG53 deficiency exacerbates DSS-induced colitis. Age-matched male WT and *MG53* knockout *(MG53*^*−/−*^) mice were administered 3% dextran sulfate sodium (DSS) in drinking water for 9 days. Body weight change (**A**) and disease activity index (DAI, **B**) were monitored daily. Representative colon images (**C**) and colon length measurements (**D**). Representative hematoxylin and eosin (H&E)-stained colon sections (**E**) and histological scores (**F**). Data are shown as mean ± SD. Statistical significance was determined by two-way ANOVA with Sidak’s multiple comparisons test (**A** and **B**) or unpaired two-tailed Student’s t-test (**D** and **F**). \*\**p* < 0.01.

Consistent with these findings, histopathological analysis using H&E staining revealed that *MG53* deficiency led to extensive inflammatory cell infiltration and more severe mucosal epithelial disruption following DSS treatment (*p* < 0.01, **Fig. 1E–F**). In WT mice, endogenous MG53 protein is normally absent in colon tissue; however, DSS challenge induced detectable MG53 accumulation in the colon, an effect absents in *MG53*^−/−^ mice (**Fig. 2A**). As we have previously shown, MG53 can be mobilized from the circulation into injured tissues, including hepatocytes or epithelial cells with low or no baseline expression of MG53, under conditions of stress such as ischemia-reperfusion or exposure to toxic agents (e.g., acetaminophen, cisplatin)(44, 45). Similarly, in the context of DSS-induced injury, MG53 was transferred from the bloodstream into the colon, serving as an emergency repair mechanism to preserve epithelial integrity.

**Figure 2.**
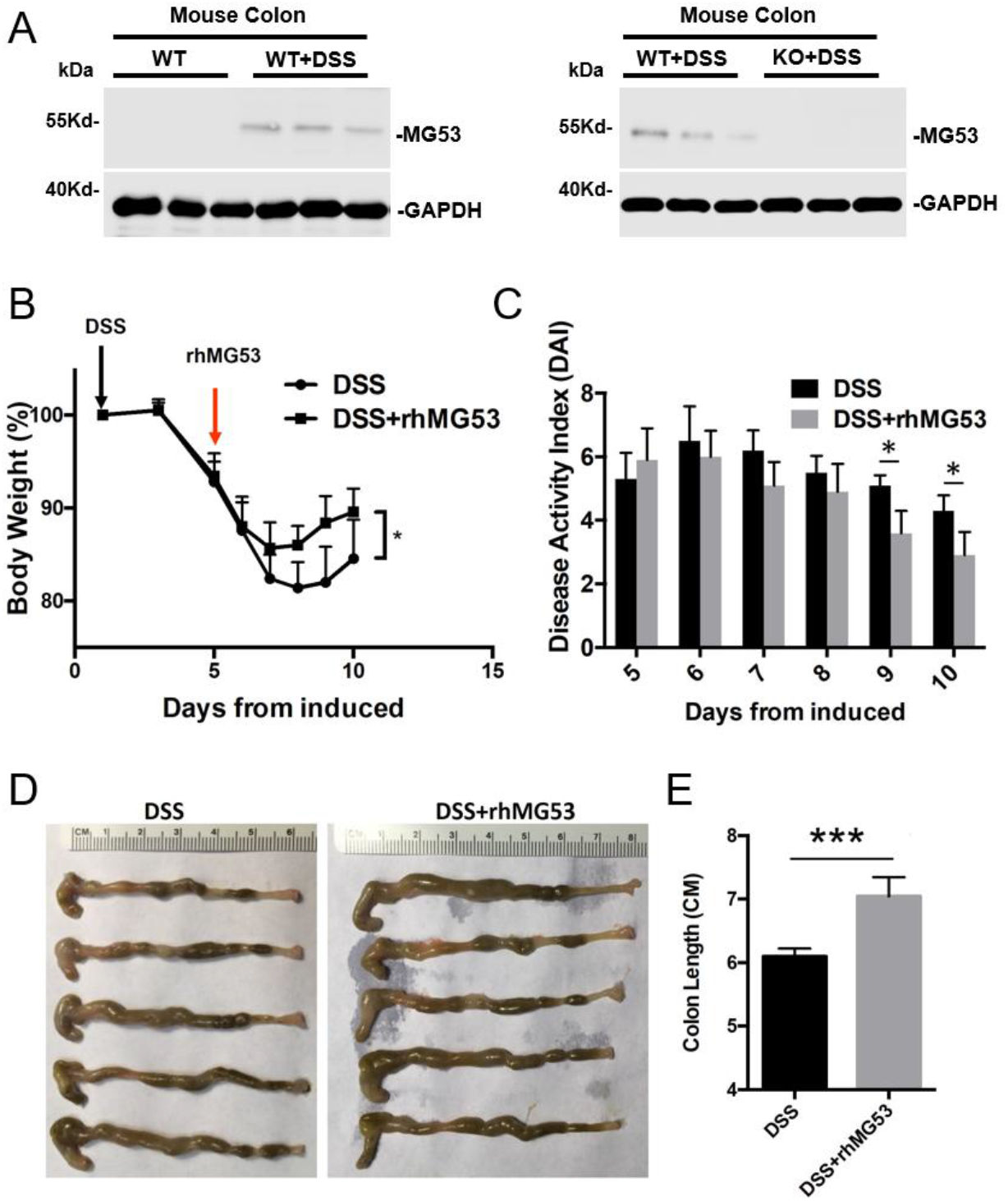
rhMG53 protein treatment mitigates DSS-induced IBD in *MG53*^-/-^ mice. *MG53*^−^/^−^ mice were administered 3% DSS in drinking water to induce colitis. Recombinant human MG53 (rhMG53, 2 mg/ml) or vehicle intraperitoneally injected daily from day 5 to day 10 (n = 5 per group). **A**. Changes in MG53 protein level in the colons from the indicated groups. **B**. Changes in body weight were monitored throughout DSS treatment. **C**. DAI scores were recorded daily. **D**. Representative images of colons collected on day 10. **E**, Quantification of colon length. Data are shown as mean ± SD; statistical significance was determined by two-way ANOVA (**B** and **C**) or Student’s t-test (**E**). Data are mean ± SD.\*\*\**p* < 0.005, \**p* < 0.05.

To establish the therapeutic potential of MG53 application, *MG53*^−/−^mice received intraperitoneal injections of rhMG53 (2mg/kg) daily from day 5 through day 10 post-DSS treatment (**Fig. 2**). Remarkably, rhMG53 administration substantially alleviated DSS-induced colitis severity. Specifically, treated mice exhibited attenuated weight loss, with separation from the DSS group emerging on day 7, followed by an earlier onset of weight recovery beginning at day 8 (*p* < 0.05, **Fig. 2B**). In line with these observation, DAI scores were significantly lower in the rhMG53-treated group compared with controls at day 9 and 10 (*p* < 0.05, **Fig. 2C**). Moreover, rhMG53 preserved colon length (*p* < 0.005, **Fig. 2D and 2E**).

These findings provide quantitative evidence for MG53’s protective function in mitigating DSS-induced colonic inflammation and demonstrate that exogenous rhMG53 can effectively rescue the exacerbated phenotype observed in *MG53*^−/−^ mice. Collectively, these data demonstrate that MG53 serves as a protective agent in maintaining integrity during colonic injury and suggests promising therapeutic potential in inflammatory bowel diseases.

### MG53 Suppresses DSS-induced Inflammatory Cytokine Release

To explore mechanisms underlying MG53’s protective effect, we investigated immune cell infiltration of the colon. Immunohistochemical analysis for CD11b, a marker of activated myeloid cells, showed a striking accumulation of CD11b-positive inflammatory cells in the colons of *MG53*^−/−^ mice compared with WT controls (*p* < 0.01, **Fig. 3A and B**). This result demonstrates *MG53* deficiency may promote excessive immune cell recruitment to site(s) of mucosal damage.

**Figure 3.**
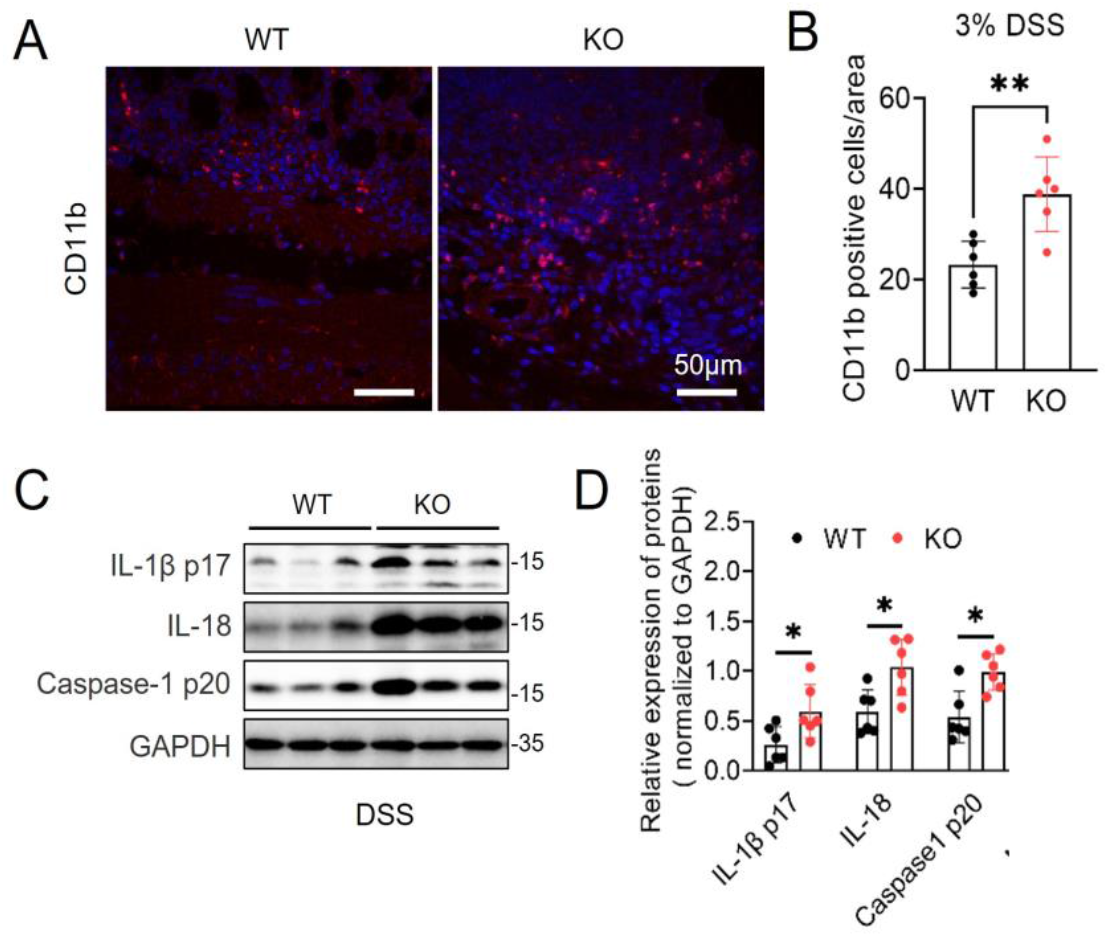
MG53 inhibits DSS-induced inflammation. **A**. Representative images of immunofluorescence CD11b (red) and DAPI (blue) in WT and *MG53*^−/−^ mice on day 9 after DSS treatment. Scale bar, 50 μm. **B**. Quantification of CD11b in panel (**A**), n = 6 for each group. **C** and **D**. Immunoblot (**C**) and quantification analysis (**D**) of mature interleukin-1β (IL-1β p17), interleukin-18 (IL-18) and cleaved caspase-1 (p20), in colon from WT and *MG53*^−/−^ mice on day 9 after DSS treatment. n = 6 each group. Data are shown as mean ± SD. Statistical significance was determined by unpaired two-tailed Student’s t-test. \*\**p* < 0.01, \**p* < 0.05.

Notably, the absence of *MG53* resulted in hyperactivation of the NLRP3 inflammasome in colon, as evidenced by substantially elevated levels of cleaved caspase-1 (p20) and markedly increased production of pro-inflammatory cytokines IL-1β and IL-18 (*p* < 0.05, **Fig. 3C and D**). Together, these results demonstrate that *MG53* deficiency exacerbates colonic inflammation by promoting immune cell infiltration and amplifying inflammasome activation, suggesting that MG53 serves as a negative regulator of colonic inflammation.

### MG53 Suppresses NLRP3 Inflammasome Formation and Attenuates Pro-inflammatory Cytokine Release

Emerging evidence has established NLRP3, a NOD-like receptor family member, as a central pathogenic driver in inflammatory bowel disease (8, 9). As a critical component of the inflammasome, NLRP3 is ubiquitously expressed across epithelial and immune cell populations, positioning it as a key therapeutic target. To investigate whether rhMG53 regulates inflammasome activation, LPS-primed bone BMDMs were treated with increasing concentrations of rhMG53 (1, 5, or 10 µg/ml) prior to nigericin stimulation.

To investigate whether rhMG53 regulates inflammasome activation, we first assessed basal MG53 expression and found that endogenous MG53 protein was undetectable in cell lysates from resting or LPS-primed WT BMDMs (Fig. 4A). Western blot analysis of culture supernatants revealed a dose-dependent reduction in both IL-1β p17 and active caspase-1 fragment (caspase-1 p20) (**Fig. 4A**), ELISA further confirmed that IL-1β secretion was significantly decreased with increasing rhMG53 concentrations (*p* < 0.05, **Fig. 4B**). To validate the broad anti-inflammasome activity of MG53, BMDMs were stimulated with ATP or MSU crystals, two additional NLRP3 inflammasome activators, and treatment with 10 µg/ml rhMG53 markedly suppressed IL-1β release and caspase-1 activation **(Fig.4C**). Consistent ELISA data demonstrated that rhMG53 significantly reduced ATP- or MSU-induced IL-1β secretion (*p* < 0.01, **Fig. 4D**). In contrast, rhMG53 exerted no detectable influence on the priming phase, as administration either preceding or following LPS stimulation failed to modify expression levels of NLRP3, ASC, pro-IL-1β, or pro-caspase-1 (**Fig. 4E**), indicating that MG53 selectively targets activation of the inflammasome signaling cascade.

**Figure 4.**
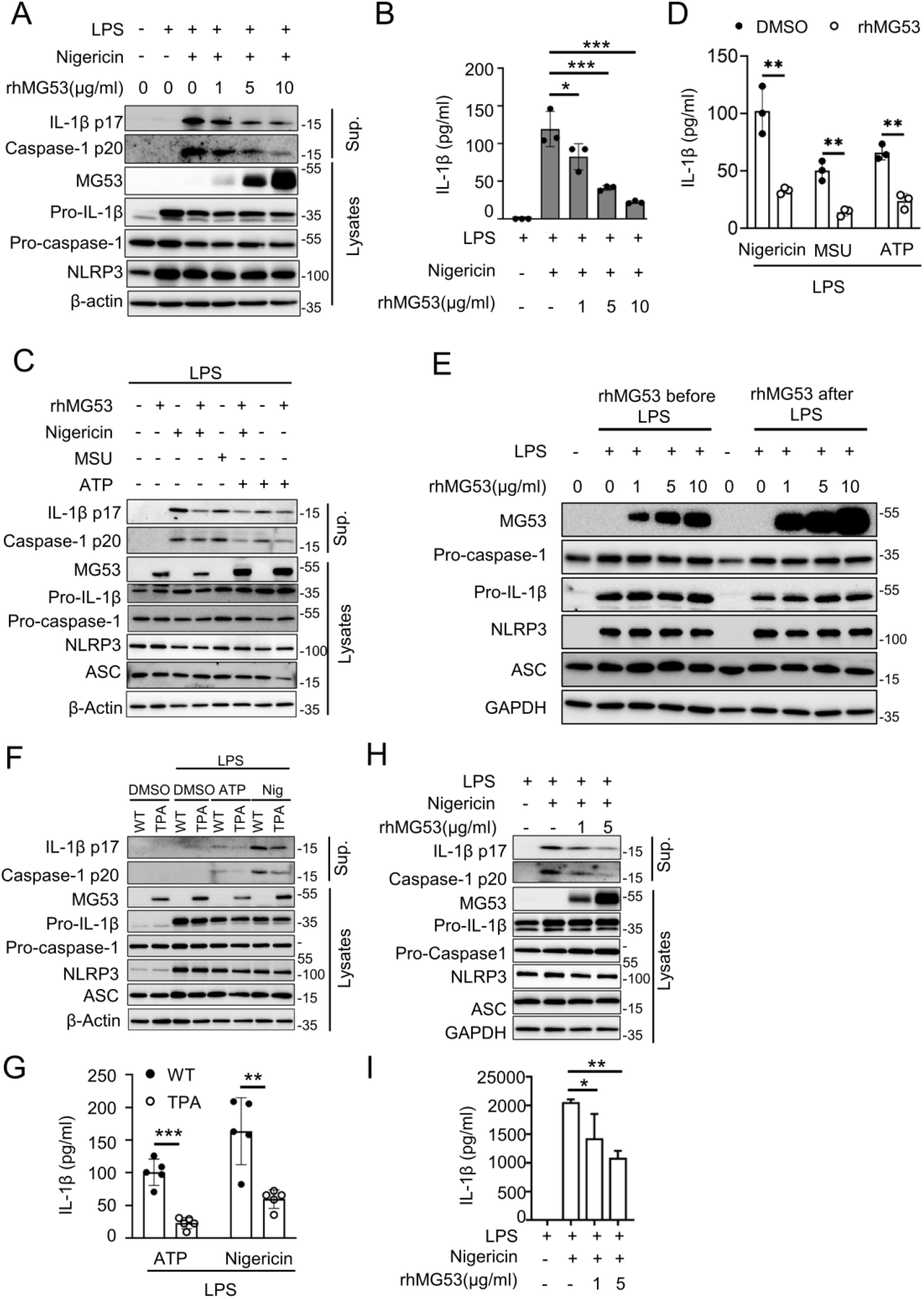
rhMG53 inhibits the activation of NLRP3 inflammasome. **A**. Immunoblot of IL-1β p17 and caspase-1 p20 in supernatants (Sup.) from lipopolysaccharide (LPS)-primed mouse bone marrow–derived macrophages (BMDMs) (100 ng/ml, 3 h) treated with rhMG53 for 1 h and stimulated with nigericin for 1 h. Pro–IL-1β, pro–caspase-1, NOD-like receptor family pyrin domain containing 3 (NLRP3), and MG53 were analyzed in cell lysates. **B**. IL-1β in culture supernatant was measured by enzyme-linked immunosorbent assay (ELISA). Treatments as in (**A**). n=3. **C**. Immunoblot of IL-1β p17 and caspase-1 p20 in supernatants from LPS-primed BMDMs treated with rhMG53 (10 μg/ml, 3 h) and stimulated with nigericin (10 μg/ml, 1 h), adenosine 5′-triphosphate disodium salt (ATP) (5 mM, 30 min), or monosodium urate crystals (MSU) (150 μg/ml, 5 h). Pro–IL-1β, pro– caspase-1, NLRP3, apoptosis-associated speck-like protein containing a CARD (ASC) and MG53 were assessed in lysates. **D**. ELISA of IL-1β release in culture supernatants. Treatments as in (**C**). n=3. E. BMDMs were treated with rhMG53 (1, 5, 10 μg/ml) for 1h before or after LPS stimulation, and NLRP3 inflammasome-related proteins were examined by immunoblotting. **F**. Immunoblot of IL-1β p17 and caspase-1 p20 in culture supernatants of LPS-primed WT or MG53-overexpressing transgenic mice (TPA) BMDMs treated for 4 h and stimulated with nigericin (10 μg/ml, 1 h) or ATP (5 mM, 30 min). MG53, NLRP3, ASC, pro-IL-1β, and pro-caspase-1 were analyzed in cell lysates. **G**. ELISA of IL-1β release in supernatants from LPS-primed WT or TPA BMDMs treated for 3 h and stimulated with nigericin (10 μg/ml,1h) or ATP (5 mM, 30 min). n=5. **H**. THP-1 cells differentiated with PMA were primed with LPS (0.5 μg/ml, 4 h), treated with rhMG53 at the indicated concentrations for 1 h, and stimulated with nigericin (1 μg/ml, 2 h). Supernatants and cell lysates were analyzed by immunoblotting. IL-1β p17 and caspase-1 p20 were detected in supernatants, while pro-caspase-1, pro-IL-1β, NLRP3, ASC, and MG53 were detected in lysates. **I**. ELISA measurement of IL-1β release in culture supernatants. n=3. Data are mean ± SD. Statistical significance was determined by one-way ANOVA (**B and I**) and unpaired Student’s t-test (**D and G**), \*\*\**p* < 0.005, \*\**p* < 0.01, \**p* < 0.05.

To further substantiate these pharmacologic observations through genetic validation, BMDMs isolated from WT or TPA mice were primed with LPS and stimulated with ATP or nigericin. Immunoblots of culture supernatants demonstrated reduced IL-1β p17 and caspase-1 p20 release in MG53-overexpressing BMDMs, while cell lysates exhibited comparable levels of NLRP3, pro-caspase-1, and pro-IL-1β (**Fig. 4F**). ELISA validation further confirmed significantly decreased IL-1β secretion in MG53-overexpressing BMDMs compared with WT controls (*p* < 0.01, **Fig. 4G**).

Additionally, these mechanistic insights were recapitulated in immortalized human monocytes. THP-1 cells differentiated into adherent macrophages with PMA treatment underwent sequential LPS pre-stimulation, rhMG53 treatment, and nigericin challenge. Western blot analysis demonstrated that rhMG53 robustly suppressed caspase-1 activation and mature IL-1β release in a concentration-dependent manner (**Fig. 4H**), with ELISA measurements providing corroborative evidence of dose-dependent IL-1β secretion inhibition (*p* < 0.05, **Fig. 4I**). Collectively, these results indicate that MG53 functions as a negative regulator of the NLRP3 inflammasome by specifically suppressing caspase -1 activation and IL-1β maturation.

### MG53 Disrupts ASC Oligomerization Interacting with NLRP3

LPS-primed BMDMs exhibited canonical recruitment of the adaptor protein ASC to NLRP3 upon nigericin stimulation, leading to the formation of a single ASC speck per cell, as visualized by confocal microscopy (**Fig. 5A**). Under control conditions, approximately 80% of cells developed ASC specks following nigericin stimulation. However, rhMG53 treatment suppressed ASC speck formation in a dose-dependent manner, reducing the proportion of speck-positive cells to ∼63%, 53%, and 34% at 1, 5, and 10 µg/ml, respectively (*p* < 0.01, **Fig. 5A and B**). Morphological evidence of inflammasome inhibition was corroborated by biochemical crosslinking assays, showing that rhMG53 markedly reduced nigericin-induced ASC oligomerization, a pivotal step in inflammasome assembly and activation. (**Fig. 5C**). Immunofluorescence staining further demonstrated that in LPS-primed BMDMs, NLRP3 was diffusely distributed in the cytoplasm and nucleus, while MG53 localized to the plasma membrane under basal conditions. Nigericin stimulation triggered dramatic NLRP3 puncta formation, which was diminished in cells treated with rhMG53. Finally, MG53 demonstrated striking colocalization with NLRP3 under combined LPS/nigericin/rhMG53 treatment conditions (**Fig. 5D and E**), suggesting spatial proximity during inflammasome modulation.

**Figure 5.**
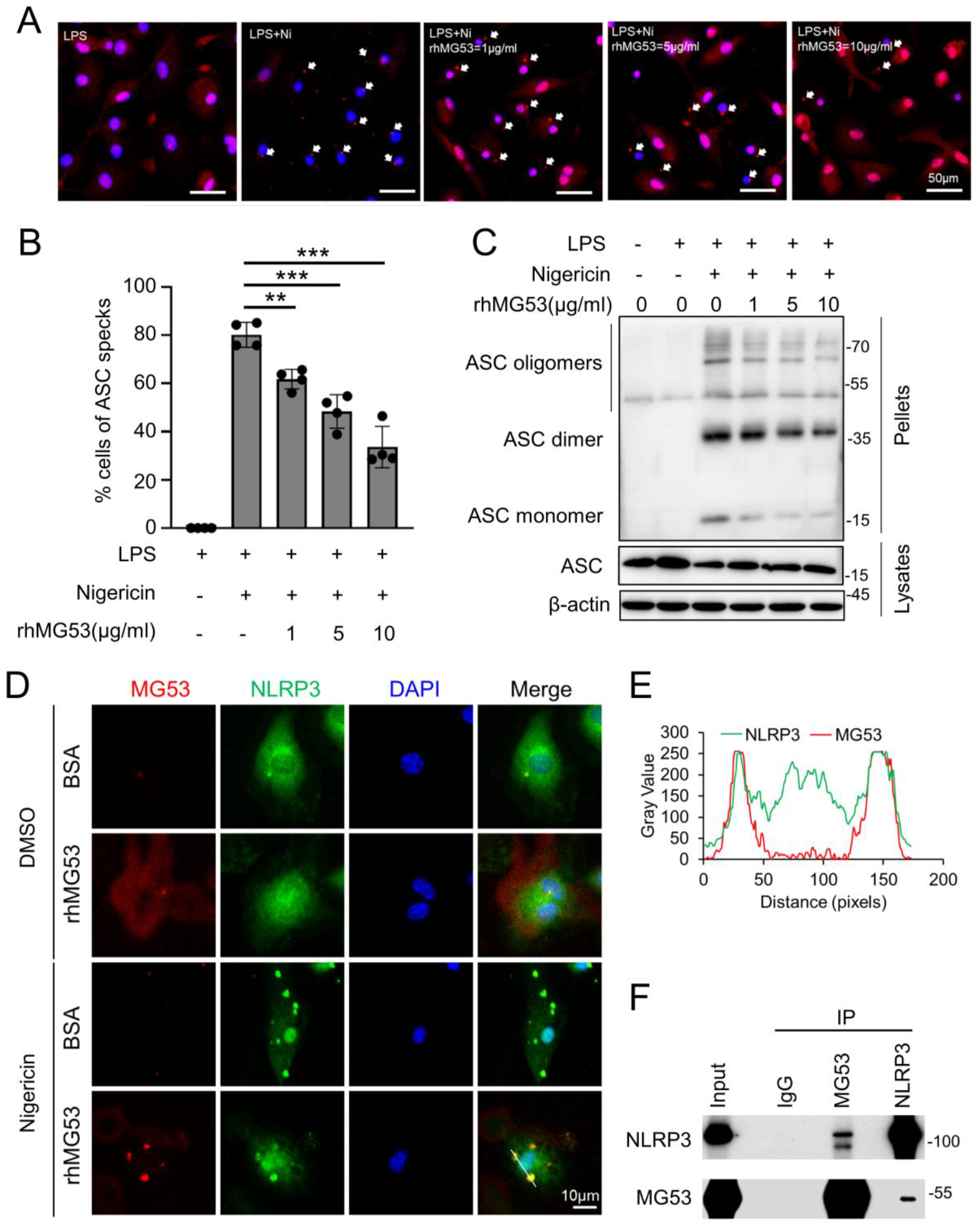
rhMG53 inhibits the formation of ASC oligomers mediated by NLRP3 inflammasome. **A**. Immunofluorescence images of ASC in BMDMs primed with LPS (100 ng/ml, 3 h), treated with rhMG53 (1, 5, 10 µg/ml) for 1 h, and stimulated with nigericin (10 μg/ml, 1 h). Nuclei were stained with DAPI (blue); ASC is shown in red. White arrows indicate ASC specks. n = 4. **B**. Quantification of ASC speck-positive cells under the conditions described in (**A**). Data are mean ± SD. \*\*\**p* < 0.005, \**p* < 0.05. **C**. ASC oligomerization and redistribution assay in BMDMs treated as in (**A**). Immunoblot analysis of ASC in DSS-crosslinked pellets (upper) and corresponding cell lysates (lower). **D**. Representative confocal images showing colocalization of NLRP3 (green) and MG53 (red), with nuclei counterstained by DAPI (blue). **E**. Fluorescence intensity profiles along the white line indicated in (**D**) revealed partial overlap of NLRP3 and MG53 signals. **F**. Immunoprecipitation was performed in LPS/nigericin- and rhMG53-treated BMDMs.

To further investigate the molecular mechanism, Co-IP assays were employed to investigate potential protein-protein associations between MG53 and NLRP3. While control IgG failed to pull down either protein, immunoprecipitation with anti-MG53 antibody successfully co-precipitated NLRP3, and reciprocally, immunoprecipitation with anti-NLRP3 antibody co-precipitated MG53 (**Fig. 5F**), providing evidence of physical interaction within the same molecular complex. Together, these findings establish a novel mechanistic paradigm whereby MG53 inhibits NLRP3 inflammasome activation. Through specific protein-protein interaction with NLRP3, MG53 disrupts the critical oligomerization of ASC adaptor proteins, thereby inhibiting preventing inflammasome assembly and subsequent caspase-1 activation during inflammasome activation.

### MG53 interacts with NLRP3 to suppress oligomerization-dependent inflammasome assembly

To elucidate the molecular dynamics of MG53-NLRP3 interaction, MG53-mCherry and NLRP3-GFP constructs were co-transfected into HEK293T cells for live-cell imaging analysis. Confocal microscopy revealed that under basal conditions, NLRP3-GFP exhibited diffuse cytoplasmic distribution. However, upon nigericin stimulation, NLRP3 underwent dramatic redistribution, forming multiple discrete punctate structures (**Fig. 6A**). Importantly, these puncta were morphologically distinct from the singular, large ASC specks observed in macrophages, suggesting that NLRP3 initially assembles into smaller oligomeric intermediates prior to recruitment into mature inflammasome complexes.

**Figure 6.**
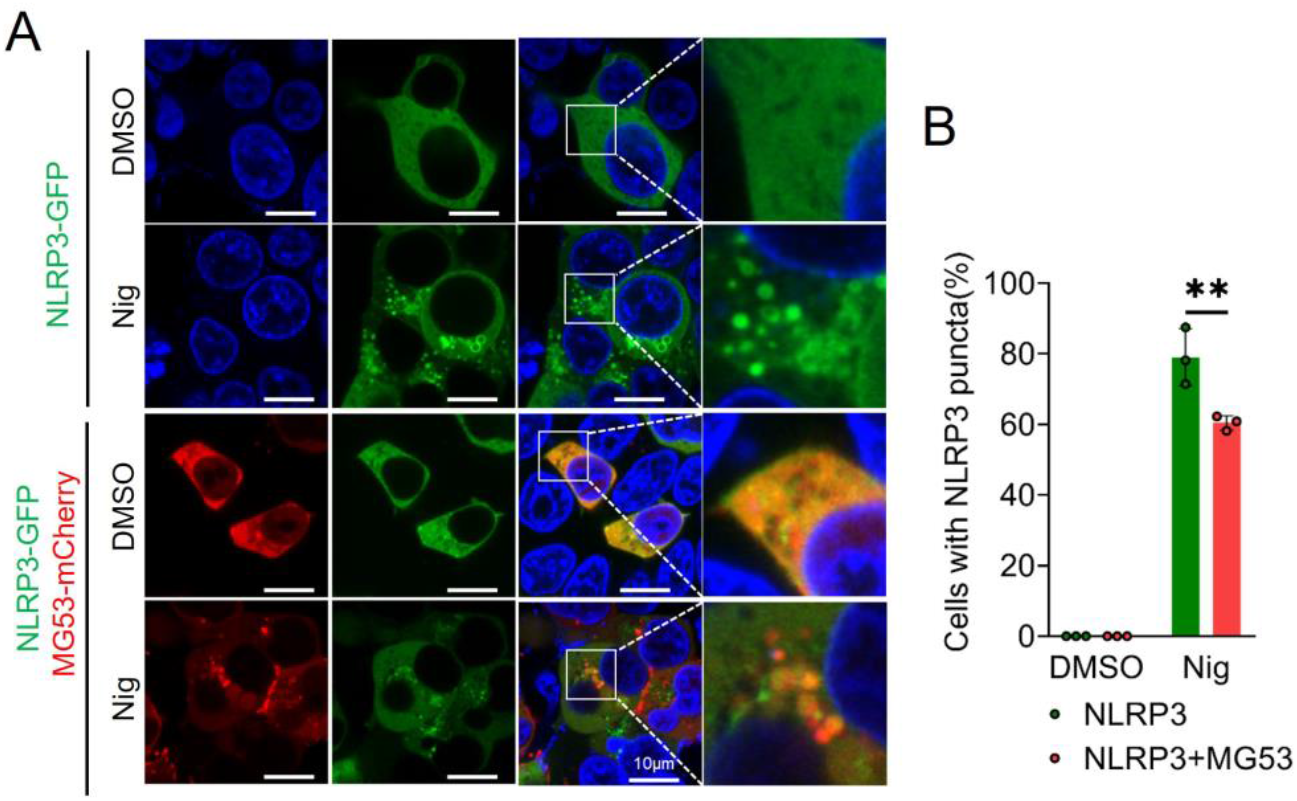
MG53 inhibits the formation of NLRP3 puncta by interacting with NLRP3. **A**. HEK293T cells were transfected with indicated plasmids for 24 h, followed by stimulation with or without 10 μM nigericin for 90 min prior to imaging. Colocalization of MG53 (red) and NLRP3 (green) was assessed by confocal microscopy. Nuclei were stained with Hoechst 33342 (blue). **B**. Quantification of the percentage of cells containing NLRP3 puncta in **A** (n = 3 repeat experiments, mean ± SD, two-sided t-test, \*\**p* < 0.01).

Co-expression of MG53-mCherry and NLRP3-GFP showed robust colocalization under both basal and nigericin-stimulated conditions, demonstrating constitutive protein-protein interaction. Moreover, nigericin-induced NLRP3 puncta was markedly diminished in cells co-expressing MG53 compared to cells expressing NLRP3 alone (*p* < 0.01, **Fig. 6B**). Collectively, these findings posit that MG53 functions as a molecular inhibitor of NLRP3 oligomerization through constitutive protein-protein interaction. This mechanism represents a novel paradigm for inflammasome regulation, positioning MG53 as a key endogenous suppressor that maintains inflammatory homeostasis through inhibition of NLRP3 assembly dynamics.

## DISCUSSION

This study uncovers a previously unrecognized role for MG53 as a critical suppressor of intestinal inflammation via inhibition of NLRP3 inflammasome activation. While MG53 has been extensively characterized as a membrane repair protein in striated muscle(19, 31, 40, 46, 47), our findings significantly expand its functional repertoire by establishing it as a key modulator of innate immune signaling and intestinal homeostasis.

Using a well-established murine model of DSS-induced colitis, we demonstrate that genetic ablation of MG53 exacerbates colonic inflammation, leading to greater weight loss, increased disease activity, shortened colon length, and pronounced histopathological damage. These findings clearly establish endogenous MG53 as a protective factor in the pathogenesis of experimental colitis. Notably, MG53 is not constitutively expressed in the colon under homeostatic conditions; however, DSS-induced injury triggers its accumulation in colonic tissue, suggesting a stress-responsive mobilization of MG53 from the circulation, consistent with prior reports in models of hepatic and renal injury(21, 44, 45, 48). This inducible recruitment may represent a generalized physiologic damage-response mechanism across epithelial organs.

Importantly, administration of rhMG53 protein significantly ameliorated colitis severity in *MG53*^−/−^ mice, restoring colon length, reducing inflammatory scores, and improving clinical outcomes. These therapeutic effects, together with the exacerbated phenotype in MG53-deficient animals, provide compelling evidence that MG53 plays an important role in preserving mucosal integrity during colonic inflammation.

Mechanistically, we uncover a novel function of MG53 as a molecular inhibitor of the NLRP3 inflammasome, a cytosolic multiprotein complex known to drive excessive cytokine release in IBD and other chronic inflammatory disorders. Our data demonstrate that MG53 inhibits NLRP3 inflammasome assembly by physically interacting with NLRP3, thereby blocking ASC oligomerization and downstream caspase-1 activation. Imaging studies revealed striking co-localization between MG53 and NLRP3 during inflammasome activation, reinforcing the concept of MG53 as a molecular brake on the early stages of NLRP3 oligomerization. This mode of inhibition is mechanistically distinct from other known negative regulators that primarily target the priming phase or promote degradation of inflammasome components. Definitive biochemical validation using purified proteins (e.g., GST pull-down or SPR) will be pursued in future studies.

The broad suppression of IL-1β release and caspase-1 activation by rhMG53 across multiple inflammasome activation stimuli, including nigericin, ATP, and MSU crystals, suggests that MG53’s effects are not stimulus-specific, but rather target the core assembly machinery of the NLRP3 inflammasome. Consistent findings in both murine BMDMs and human THP-1 macrophages highlight the translational potential of MG53-based interventions. The ability of exogenous rhMG53 to rescue the heightened inflammatory phenotype in knockout mice further underscores its promise as a viable therapeutic candidate for IBD, particularly in individuals with impaired or insufficient endogenous MG53 responses.

Our work also adds to the growing appreciation of MG53’s broader physiological roles outside of striated muscle. Recent studies have implicated MG53 in epithelial repair, stem cell differentiation, and mitochondrial protection(49, 50). Notably, *Pei et al*. recently described an MG53-PPARα axis that promotes secretory lineage differentiation in intestinal stem cells, further supporting the relevance of MG53 in gastrointestinal health(50). Distinct from this cell-autonomous mechanism, our data support circulating MG53 is enriched in the inflamed colon and taken up by infiltrating macrophages to inhibit the NLRP3 inflammasome. Thus, our study complements the regenerative findings of Pei *et al*. by defining a macrophage-dependent paracrine pathway that operates in concert with epithelial repair. Future research requires the adoption of cell type-specific genetic models and lineage tracing strategies to thoroughly elucidate the mechanisms of the MG53 signaling pathway in epithelial and immune tissues. Taken together, these findings position MG53 as a multifaceted regulator of epithelial resilience and immune modulation, with applications extending beyond the context of colitis.

Despite these advances, several key questions remain. CD11b is a pan-myeloid marker. While its reduction confirms broad inflammatory suppression, it does not distinguish between macrophages and neutrophils. Given that macrophages drive this cascade via NLRP3, we focused on BMDMs for mechanistic insight. Future investigations using lineage-specific markers (e.g., F4/80, Ly6G) will further characterize the dynamics of these distinct immune populations. Although rhMG53 enters macrophages to inhibit NLRP3 inflammasome activation, the precise mechanism of its internalization remains to be elucidated. The precise domains mediating MG53-NLRP3 interaction have yet to be mapped. Defining these interfaces may guide the development of targeted mimetics or small molecules that replicate MG53’s anti-inflammatory effects. Furthermore, the regulation of MG53 expression and activity in immune and epithelial cells under physiological and pathological conditions remains largely unexplored. It also remains to be determined whether MG53 modulates other inflammasomes (e.g., AIM2, NLRC4) or intersects with additional innate immune pathways such as STING or NF-*κ*B signaling.From a translational standpoint, it will be important to assess MG53 expression levels and functional variants in human IBD patients, as deficiency or dysfunction may predispose individuals to heightened inflammasome activation and more severe disease. Additionally, expanding these studies to other colitis models (e.g., TNBS, adoptive T-cell transfer, IL-10 knockout) and ultimately to clinical trials will be critical steps toward establishing the therapeutic utility of MG53 in human IBD.

The NLRP3 inflammasome is a known driver of pathogenesis in numerous chronic diseases, including atherosclerosis, neurodegeneration, metabolic syndrome, and fibrotic disorders(46). Thus, MG53-based strategies may offer broad utility in combating diverse NLRP3-driven pathologies, potentially opening a new therapeutic class of inflammasome-targeted protein biologics.

## Conclusion

In summary, this study identifies MG53 as an endogenous inhibitor of NLRP3 inflammasome activation and establishes its critical role in protecting against experimental colitis. Through interaction with NLRP3, MG53 disrupts inflammasome assembly, inhibits caspase-1 activation, and attenuates pro-inflammatory cytokine release. These findings reveal a previously unappreciated dimension of MG53 function in immune regulation and highlight its potential as a therapeutic agent for IBD and other inflammasome-mediated diseases. The efficacy of exogenous rhMG53 in rescuing severe colitis provides a strong foundation for future translational development.

## Acknowledgement

We acknowledge the core facilities for microscopy, flow cytometry, and animal care that made this work possible. This work was partially supported by NIH grants to J.M. (R01HL157215, R01AG071676, R01AG072434, R01EY036243), H.L. (R01AG056919, R21AR080628), and the OSU Comprehensive Cancer Center Startup Fund to H.L.

## Data Availability Statement

The datasets for this study are available from the corresponding author on reasonable requests.

## Conflict of Interest

JM is a founder for TRIM-edicine, Inc, which develop the MG53 technology for regenerative medicine application. All other authors declare no competing financial interests, professional affiliations, or personal relationships that could have influenced the conduct, analysis, or reporting of the research presented in this manuscript.

